# Macaques preferentially attend to intermediately surprising information

**DOI:** 10.1101/2021.08.02.454786

**Authors:** Shengyi Wu, Tommy Blanchard, Emily Meschke, Richard N. Aslin, Benjamin Y. Hayden, Celeste Kidd

**Affiliations:** Department of Psychology, University of California, Berkeley, 2121 Berkeley Way West, Berkeley, CA 94720; Klaviyo, 125 Summer St, Floor 6, Boston, MA 02111; Helen Wills Neuroscience Institute, University of California, Berkeley, 175 Li Ka Shing Center, MC#3370, Berkeley, CA 94720; Haskins Laboratories, Yale University, 300 George Street, New Haven, CT 06511; Department of Neuroscience, University of Minnesota, 321 Church St SE, Minneapolis, MN 55455

**Keywords:** Attention, statistical learning, eye tracking

## Abstract

Normative learning theories dictate that we should preferentially attend to informative sources, but only up to the point that our limited learning systems can process their content. Humans, including infants, show this predicted strategic deployment of attention. Here we demonstrate that rhesus monkeys, much like humans, attend to events of moderate surprisingness over both more and less surprising events. They do this in the absence of any specific goal or contingent reward, indicating that the behavioral pattern is spontaneous. We suggest this U-shaped attentional preference represents an evolutionarily preserved strategy for guiding intelligent organisms toward material that is maximally useful for learning.

## Introduction

Intelligent organisms acquire knowledge through experience; however, there is more information available than they can actually explore [1, 2]. Thus, intelligent organisms must be selective.

Adaptive theories of curiosity posit that uncertainty helps guide learners’ exploration [3, 4, 5, 6, 7, 8, 9]. Specifically, adaptive learners attend to information of intermediate uncertainty. This results in a U-shaped relationship between uncertainty and inattention: Low uncertainty events offer little to learn from and high uncertainty events are beyond the learners’ processing capabilities [3, 8, 10, 11, 12, 13, 14,15,16,17,18]. This mechanism has been attested in humans, and may represent an elegant solution for intelligent organisms to resolve the information overload problem.

Human infants and children preferentially maintain attention to sequential events of intermediate surprisal values [16, 17, 18, 19]. While this pattern has not been observed in non-humans, monkeys can seek information for its inherent value. For example, macaques will sacrifice some liquid reward in exchange for information with no strategic benefit [20, 21] and engage in directed exploration [22, 23]. These data raise the possibility that strategic information-seeking patterns may reflect an evolutionarily ancient capacity for adaptive regulation of incoming information. If so, this would demonstrate a general principle of advanced evolved learners rather than a uniquely human skill.

Here, we employ a variation on the infant paradigm with rhesus macaques. We test the hypothesis that adaptive regulation of information-seeking is a cognitive skill shared with our common ancestor. Unlike most previous work on curiosity in macaques, we employ a free-viewing paradigm without rewards tied to particular responses. This approach tests for spontaneous preference and avoids possible learning effects. We find that macaques’ visual attention is strikingly similar to that of human infants.

## Methods

### Subjects

All animal procedures were performed at the University of Rochester (Rochester, NY, USA) and were approved by the University of Rochester Animal Care and Use Committee. Experiments were conducted in compliance with the Public Health Service’s Guide for the Care and Use of Animals. Five male rhesus macaques (*Macaca mulatta*) served as subjects^1^. Subjects had full access to standard chow in their home cages. Subjects received at minimum 20 mL/kg water per day, although in practice they received close to double this amount in the lab as a result of our experiments. Subjects had been trained to perform oculomotor tasks for liquid rewards through positive-reward-only reinforcement training.

### Stimuli

Visual stimuli were colored shapes on a computer monitor (see Figure 1a). We designed the displayed stimuli to be easily captured by a simple statistical model [16, 18]. Each trial featured one of 80 possible visual-event sequences (See Appendix 1). All sequences were presented to all subjects in different randomized orders. One sequence was presented per trial, and each was presented in the form of a unique animated display. An example video can be seen at haydenlab.com/surprisal.

**Figure 1:**
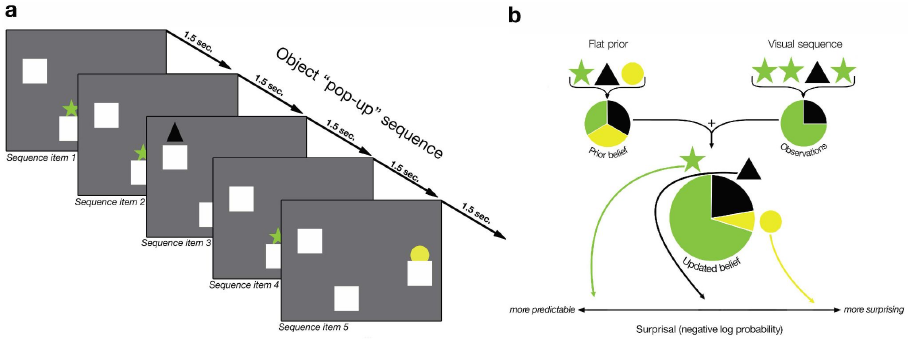
**a) Example of Sequential Visual Display.** The illustration shows five different time-points in the sequence. At each event in the sequence, one of the three unique objects popped up from behind one of three boxes. **b) Idealized Learning Model Schematic**. The schematic shows an example of how the idealized learning model forms probabilistic expectations about the expectedness of the next event in a sequence.

Each animated display featured three identical boxes in three distinct, randomly chosen spatial locations on the screen that remained static throughout the sequence. Each box concealed one unique geometric object, which was randomly selected from a set that included 4 different shapes in 8 colors (e.g., a yellow triangle, a red star, or a blue circle). Geometric objects remained associated with their respective gray boxes throughout the sequence and were each unique within a trial, but were chosen randomly from the set across trials [16, 17, 18, 19].

Objects appeared from boxes on the displays according to the sequence orders. Each *event* within a sequence consisted of one of the three unique objects popping out from behind one of the three boxes (750 ms), and then back into the box (750 ms) without overlap or delay. The 80 sequences were generated to maximize the difference of their theoretical information property, such that the pop-up probabilities of each geometric object varied if a different sequence was observed. For example, if a sequence starts with 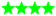 and follows by another 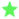, this is an example of a very predictable sequential event (low surprisal). If the same sequence starts again with 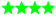 but follows by ▴, this would be an example of a less predictable sequential event (high surprisal).

### Procedure

We recorded eye movements as subjects watched the visual displays. Eye positions were measured with the Eyelink Toolbox and were sampled at 1000 Hz by an infrared eye-monitoring camera system (SR Research, Osgoode, ON, Canada) [24]. A solenoid valve delivered a 53 µL water reward when each object was at its peak (every 1.5 sec.), regardless of where or whether the subject was looking. The intermittent and fully predictable reward is a standard procedure in primate behavior studies designed to increase general task participation and arousal without making any particular task events reward-associated [25]. Regardless of subjects’ gaze behavior, each sequence (one per trial) was displayed in full. The rate of presentation was between 0 and 2 trials, interspersed within between 400 and 2,000 unrelated trials for other studies [25, 26, 27]. We recorded all spatial and temporal details of the randomized, Matlab-generated sequential displays we presented, as well as all macaque visual fixations during the stimulus presentations.

### Analysis

We analyzed three behavioral measures: reaction time, predictive-looking, and look-away. ***Reaction time (RT)*** measures the latency to shift gaze to the object after it appears. This is a standard measure to detect agents’ expectations. ***Predictive-looking***^***2***^ is a binary variable that indicates whether the subject was already looking at the current object when it first became active but before the object actually popped up. ***Look-away***^***3***^ is the first point in the sequence when the macaque looked off-screen for 0.75 sec (50% of the total pop-up event duration) [16, 17, 18]. We analyzed these three behavioral measures as a function of the surprisal value of each event in the sequence, which is the negative log probability of the event’s occurrence, according to unigram and bigram Markov Dirichlet-Multinomial (ideal observer) models (following the analysis methods of [16, 17, 18])(Appendix 6). The unigram model treats each event as statistically independent, while the bigram model assumes event order dependence and tracks the conditional probability on the immediately preceding event. The models begin with a simple prior corresponding to the implicit beliefs a learner possesses before making any observations. By using a flat (or uninformative) prior, we assume that the learner begins the sequence presentation with the implicit belief that each of the three possible objects are equally likely to pop-up from behind their occluding boxes. Once the sequence presentation begins, the model estimates the surprisal value of the current event at each item in the sequence. It combines the simple prior with the learner’s previous observations from the sequence in order to form a posterior or updated belief. The next object pop-up event then conveys the surprisal value according to the probabilistic expectations of the updated belief (see Figure 1b). We also evaluated the statistical significance of each variable using mixed effect linear and logistic regressions with random intercepts, and linear and quadratic surprisal slopes. A generalized additive model (GAM) was used to visualize the relationship between the surprisal estimate from the computational model and the behavioral data.

## Results

### Quicker deployment of gaze for events of intermediate surprisal

The unigram GAM analysis shows that the relationship between reaction times and subjects’ expectations about stimulus predictability is U-shaped, with subjects exhibiting the fastest RTs for intermediately predictable stimuli (Figure 2a.1). The regression reveals both a significant linear term (*β* = -64.68, *t* = –9.42, *p* < 0.0002) and a significant quadratic term (*β* = 14.372, *t* = 6.37, *p* < 0.002). The U-shape relationship also holds when other variables, such as whether the object is a repeat, distance between the current and previous pop up object, trial number, and item number in a sequence, are controlled in the GAM model, as well as revealed by the significant linear (*β* = -26.19, *t* = –3.02, *p* < 0.02) and quadratic (*β* = 6.00, *t* = 2.62, *p* < 0.03) terms in the controlled regression (Figure 2a.2). The significance of the quadratic term likely corresponds to a genuine U over the range of surprisal, especially in light of the fact that the significance holds even in the controlled GAM. However, in the GAM analysis for the transitional surprisal measures, it shows a much shallower U-shape, with only the linear trend being significant in the raw model. Once all predictors are included, the curve becomes mostly flat. This shows that the unigram model is more robust than the transitional model to capture the relationship between subjects’ RTs and the surprisingness of stimuli. Our results also show that all five subjects exhibit similar preference for stimuli of intermediate surprisal, suggesting that the U-shape relationship holds within rhesus macaques and is not due to subject average. This consistent pattern observed in each macaque subject was also found within individual human infants who reserve attention for events that are moderately predictable [19].

**Figure 2:**
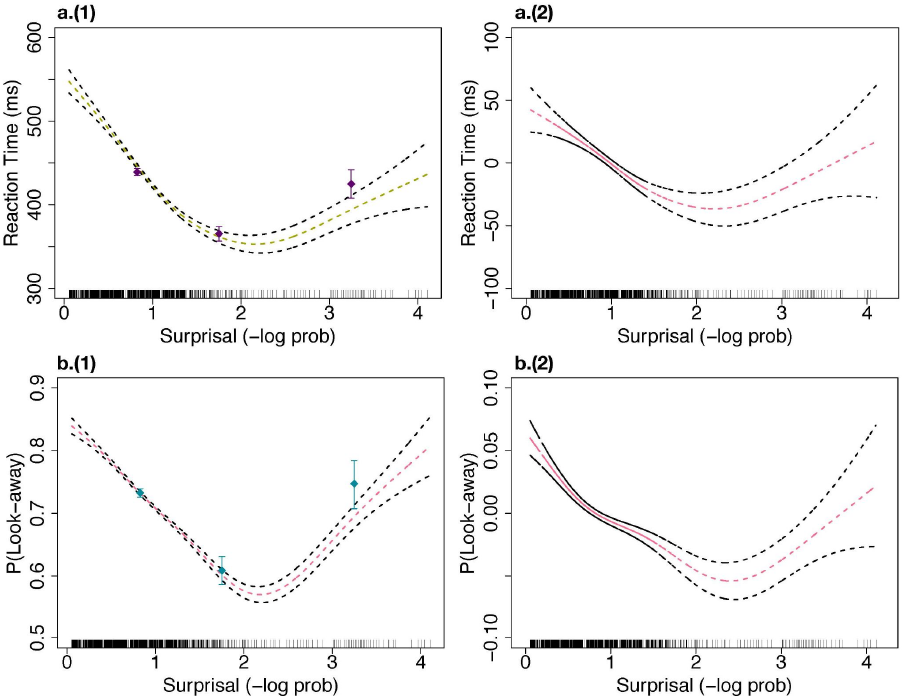
**a) Reaction Time (ms) as a Function of Unigram Surprisal.** *(1)* Subjects’ reaction time to fixate the active object (y-axis) as a function of unigram surprisal (x-axis). The points and error bars show raw data binned according to surprisal values; the smooth curve shows the fit of a generalized additive model with standard errors. Vertical tick marks show values of surprisal attained in the experiment. *(2)* Reaction Time (*y*–axis) and unigram surprisal (*x*-axis), while controlling for all factors. **b) Look-Away Probability as a Function of Unigram Surprisal**. *(1)* Subjects’ probability of looking away (y-axis) as a function of unigram surprisal (x-axis). The smooth curve shows the fit of a generalized additive model with 95% confidence interval. *(2)* The relationship between look-away probability (y–axis) and unigram surprisal (x-axis), while controlling for all covariate factors.

### Predictive looks towards unshown items

Subjects are more likely to predictively look at objects on their first appearance when the pop-up events are estimated to be more likely, according to the model. The estimated GAM plot shows a decreasing trend between the probability of predictive-looking and the surprisal value (See Appendix 2). The pattern is supported by the unigram controlled regression model, which finds statistical significance in the linear trend (*β* = -2.72, *z* = -2.65, *p* < 0.009). Subjects are also more likely to predictively look at never shown objects if they appear earlier in a sequence (*β* = -12.61, *z* = -2.13, *p* < 0.04). These results show that subjects might be curious about unknown information and spontaneously track the incoming statistics, expecting that there’s some change that will occur and, if it does, it will be informative. They also suggest that over time, as it’s increasingly unlikely to see unopened boxes ever open, macaques are less likely to allocate their attentional resources towards monitoring unopened boxes. It is further evidence that macaques’ information-seeking behavior is moderated by their expectation in the absence of overt rewards.

### Preferential gaze towards events of intermediate surprisal

Estimated by the unigram GAM analysis, subjects were more likely to terminate attention to highly predictable events and also highly unexpected events (Figure 2b.1). In the regression that considers only surprisal and squared surprisal measures, both the linear term (*β* =-0.45, *z* =-5.87, *p* < 0.001) and the quadratic term (*β* =0.11, *z* =3.679, *p* < 0.001) are statistically significant. However, the controlled logistic regression does not reveal a statistically robust quadratic term (*β* = 0.03, *z* = 0.86, *p* < 0.40). These results suggest that the U-shaped look-away attentional trend may depend upon the learner maintaining some control over what they observe. Results from the transitional model show that there is a U-shaped relationship in the raw model and model fits, with the quadratic trend being statistically significant (*β* =0.06, *z* =2.69, *p* < 0.008). However, this pattern disappears when other variables are controlled.

## Discussion

Humans do not indiscriminately absorb any information they encounter. Instead, we preferentially seek out information that is maximally useful [1, 2, 8, 28]. This regulated information-gathering strategy favors moderately surprising events, resulting in an inverse-U-shaped pattern between event surprisal and engagement. Here we show that this pattern, previously only observed in humans [16, 17, 18, 19], is also observed in rhesus macaques, a primate species that diverged from humans roughly 25 million years ago.

The presence of this pattern in macaques suggests that the capacity to adaptively seek useful information is not uniquely human, but instead reflects long-standing evolutionary pressures present since at least the time of our last common ancestor. This is important because a good deal of theorizing highlights the uniqueness of human curiosity, with the implication that curiosity is a factor that has driven human divergence [29]. Our results, then, suggest an alternative hypothesis that humans and animals share a broad suite of cognitive adaptations, and that humans differ in quantity but not in quality from non-human animals. Unlike humans [16,17], unigram statistics were more robust predictors of monkey learners’ behaviors than the transitional statistics. This difference could suggest a species-level difference; however, this conclusion is premature because the tested macaques had substantial experience with tasks for which tracking unigram statistics is more relevant (e.g., k-arm bandit tasks).

Broadly, these results highlight that just as intelligent organisms forage for food, they deliberately seek a rich and balanced diet of information that can drive maximally efficient learning and, ultimately, adaptive fitness [30].

## Acknowledgements

We thank Steven T. Piantadosi, Hayden lab, and Kidd lab for useful feedback and suggestions.

## Data Accessibility

All data and code can be found at https://github.com/shengyiwu/Monkilock

## Authors’ contributions

Conceptualization: CK, TB, RA, BH

Data curation: CK, TB

Formal analysis: SW, EM, CK

Investigation: SW, EM, CK, TB

Methodology: CK, TB, RA, BH, SW

Software: TB, BH

Visualization: SW

Writing – original draft: SW, CK

Writing – review & editing: SW, CK, RA, BH

Funding acquisition: CK, RA, BH

All authors gave final approval for publication and agree to be held accountable for the work performed therein.

## Footnotes

1 Using single-sex macaques would prevent fighting and overmating among subjects. Sex differences are not expected in macaques’ behaviors based on the results from the infant version of this study [16].

2 In this paradigm, the pop-up events occurred temporally predictably (every 1,500 ms), with no breaks between. Thus, these predictive looks differ from those elicited in most predictive-looking paradigms, where delays between events specifically encourage predictive looking. Regardless, predictive looks may be taken to indicate some degree to attentional allocation to inactive boxes in advance of their opening.

3 In this version, sequences kept unfolding no matter whether the macaques were looking. This is different from the infant paradigm, in which infants could terminate the displays with their inattention.

