## Appendix for "Macaques preferentially attend to intermediately surprising information"

### Appendix 1: Visual Sequences

Sequence Position (positions 1–30)

| Sequence ID | 1 | 2 | 3 | 4 | 5 | 6 | 7 | 8 | 9 | 10 | 11 | 12 | 13 | 14 | 15 | 16 | 17 | 18 | 19 | 20 | 21 | 22 | 23 | 24 | 25 | 26 | 27 | 28 | 29 | 30 |
|---|---|---|---|---|---|---|---|---|---|---|---|---|---|---|---|---|---|---|---|---|---|---|---|---|---|---|---|---|---|---|
| 1 | 2 | 1 | 2 | 2 | 2 | 1 | 3 | 3 | 3 | 1 | 3 | 1 | 1 | 2 | 2 | 1 | 1 | 1 | 2 | 3 | 2 | 2 | 1 | 2 | 1 | 1 | 1 | 3 | 2 | 2 |
| 2 | 3 | 3 | 3 | 3 | 1 | 3 | 3 | 1 | 3 | 3 | 3 | 2 | 3 | 3 | 3 | 1 | 1 | 1 | 3 | 3 | 1 | 3 | 1 | 2 | 1 | 1 | 1 | 2 | 2 | 2 |
| 3 | 1 | 1 | 1 | 1 | 1 | 1 | 1 | 3 | 3 | 3 | 2 | 2 | 1 | 2 | 3 | 3 | 3 | 2 | 2 | 1 | 2 | 2 | 3 | 1 | 1 | 2 | 3 | 1 | 1 | 3 |
| 4 | 3 | 2 | 1 | 2 | 2 | 1 | 2 | 1 | 1 | 3 | 2 | 2 | 1 | 1 | 2 | 2 | 2 | 1 | 1 | 1 | 2 | 1 | 1 | 1 | 2 | 2 | 1 | 2 | 2 | 2 |
| 5 | 3 | 1 | 3 | 3 | 3 | 3 | 3 | 3 | 3 | 2 | 2 | 3 | 1 | 3 | 3 | 2 | 2 | 2 | 1 | 3 | 2 | 3 | 1 | 3 | 3 | 1 | 2 | 3 | 2 | 3 |
| 6 | 1 | 1 | 1 | 1 | 1 | 1 | 1 | 1 | 2 | 2 | 2 | 1 | 2 | 3 | 3 | 1 | 2 | 2 | 1 | 2 | 3 | 2 | 3 | 3 | 2 | 2 | 1 | 3 | 3 | 1 |
| 7 | 2 | 1 | 1 | 2 | 1 | 1 | 3 | 3 | 1 | 2 | 1 | 1 | 2 | 2 | 1 | 2 | 2 | 2 | 2 | 2 | 2 | 1 | 1 | 2 | 2 | 2 | 2 | 2 | 2 | 1 |
| 8 | 1 | 1 | 1 | 1 | 1 | 3 | 1 | 1 | 2 | 3 | 3 | 1 | 1 | 1 | 3 | 1 | 1 | 1 | 2 | 3 | 2 | 1 | 3 | 3 | 2 | 2 | 3 | 2 | 2 | 2 |
| 9 | 1 | 3 | 1 | 1 | 3 | 1 | 3 | 2 | 1 | 3 | 3 | 1 | 2 | 1 | 1 | 1 | 1 | 1 | 3 | 3 | 2 | 1 | 1 | 1 | 1 | 1 | 2 | 1 | 1 | 2 |
| 10 | 2 | 1 | 1 | 3 | 2 | 3 | 3 | 2 | 1 | 3 | 2 | 3 | 3 | 1 | 1 | 1 | 1 | 3 | 1 | 3 | 1 | 1 | 3 | 3 | 3 | 1 | 3 | 3 | 1 | 3 |
| 11 | 1 | 1 | 3 | 2 | 1 | 2 | 2 | 3 | 3 | 2 | 1 | 1 | 2 | 2 | 1 | 2 | 1 | 1 | 3 | 2 | 1 | 1 | 1 | 2 | 2 | 1 | 2 | 1 | 2 | 1 |
| 12 | 1 | 2 | 1 | 2 | 3 | 1 | 2 | 2 | 3 | 1 | 2 | 1 | 2 | 3 | 2 | 2 | 2 | 2 | 2 | 1 | 1 | 3 | 3 | 3 | 2 | 3 | 3 | 1 | 3 | 2 |
| 13 | 3 | 3 | 2 | 3 | 1 | 2 | 2 | 1 | 1 | 2 | 3 | 3 | 3 | 3 | 2 | 1 | 1 | 2 | 3 | 3 | 3 | 2 | 1 | 1 | 1 | 2 | 1 | 2 | 1 | 2 |
| 14 | 3 | 2 | 2 | 3 | 3 | 2 | 2 | 2 | 3 | 3 | 2 | 3 | 3 | 2 | 2 | 3 | 3 | 2 | 3 | 2 | 2 | 2 | 3 | 2 | 3 | 3 | 1 | 1 | 1 | 3 |
| 15 | 2 | 3 | 1 | 2 | 1 | 1 | 2 | 1 | 1 | 2 | 2 | 1 | 3 | 2 | 2 | 1 | 2 | 1 | 2 | 1 | 2 | 3 | 2 | 1 | 1 | 3 | 1 | 1 | 3 | 3 |
| 16 | 2 | 2 | 2 | 2 | 2 | 1 | 2 | 2 | 1 | 2 | 1 | 1 | 2 | 1 | 1 | 1 | 1 | 2 | 3 | 2 | 2 | 2 | 2 | 3 | 3 | 3 | 1 | 3 | 3 | 1 |
| 17 | 1 | 2 | 3 | 1 | 2 | 3 | 3 | 1 | 2 | 3 | 2 | 3 | 1 | 2 | 2 | 2 | 3 | 1 | 3 | 3 | 1 | 2 | 2 | 3 | 1 | 3 | 1 | 1 | 3 | 2 |
| 18 | 2 | 1 | 2 | 1 | 1 | 2 | 1 | 1 | 1 | 1 | 3 | 2 | 2 | 2 | 2 | 2 | 3 | 2 | 1 | 2 | 2 | 2 | 3 | 2 | 3 | 1 | 2 | 3 | 3 | 2 |
| 19 | 2 | 3 | 3 | 1 | 2 | 2 | 2 | 2 | 3 | 3 | 2 | 3 | 3 | 3 | 2 | 2 | 2 | 2 | 2 | 2 | 2 | 2 | 3 | 2 | 3 | 3 | 2 | 2 | 2 | 3 |
| 20 | 1 | 3 | 3 | 1 | 3 | 3 | 2 | 2 | 1 | 2 | 2 | 3 | 2 | 2 | 3 | 1 | 1 | 2 | 2 | 3 | 1 | 1 | 2 | 2 | 3 | 2 | 1 | 2 | 2 | 2 |
| 21 | 3 | 1 | 1 | 1 | 1 | 3 | 1 | 3 | 3 | 3 | 1 | 3 | 3 | 1 | 3 | 3 | 1 | 3 | 1 | 1 | 3 | 1 | 2 | 2 | 3 | 2 | 1 | 1 | 2 | 3 |
| 22 | 2 | 1 | 2 | 2 | 1 | 3 | 1 | 2 | 2 | 2 | 2 | 1 | 3 | 2 | 2 | 2 | 2 | 2 | 1 | 3 | 2 | 2 | 2 | 1 | 3 | 2 | 1 | 2 | 1 | 3 |
| 23 | 3 | 2 | 1 | 3 | 2 | 1 | 1 | 3 | 1 | 2 | 3 | 2 | 1 | 1 | 2 | 1 | 2 | 1 | 1 | 1 | 1 | 2 | 1 | 1 | 1 | 2 | 1 | 1 | 2 | 2 |
| 24 | 2 | 1 | 1 | 1 | 3 | 3 | 1 | 3 | 1 | 1 | 3 | 3 | 3 | 1 | 3 | 3 | 1 | 3 | 2 | 1 | 1 | 2 | 2 | 3 | 1 | 2 | 1 | 1 | 2 | 2 |
| 25 | 3 | 2 | 1 | 3 | 1 | 3 | 2 | 2 | 1 | 3 | 1 | 3 | 1 | 2 | 3 | 1 | 1 | 3 | 3 | 2 | 3 | 2 | 1 | 3 | 3 | 3 | 3 | 2 | 1 | 2 |
| 26 | 3 | 3 | 3 | 3 | 3 | 3 | 3 | 2 | 2 | 1 | 2 | 1 | 2 | 1 | 1 | 1 | 2 | 3 | 1 | 1 | 3 | 1 | 3 | 1 | 2 | 1 | 3 | 3 | 2 | 2 |
| 27 | 2 | 2 | 1 | 2 | 3 | 1 | 1 | 3 | 2 | 2 | 3 | 3 | 2 | 2 | 3 | 1 | 1 | 3 | 1 | 3 | 1 | 1 | 3 | 1 | 1 | 1 | 1 | 1 | 3 | 2 |
| 28 | 2 | 3 | 1 | 3 | 3 | 1 | 1 | 1 | 2 | 2 | 3 | 3 | 1 | 3 | 3 | 1 | 2 | 2 | 3 | 2 | 2 | 3 | 3 | 2 | 1 | 2 | 1 | 3 | 3 | 2 |
| 29 | 1 | 1 | 1 | 1 | 1 | 1 | 1 | 1 | 1 | 1 | 1 | 1 | 1 | 1 | 1 | 1 | 1 | 1 | 1 | 1 | 1 | 1 | 1 | 1 | 1 | 1 | 1 | 1 | 1 | 1 |
| 30 | 1 | 2 | 1 | 2 | 1 | 2 | 1 | 2 | 1 | 2 | 1 | 2 | 1 | 2 | 1 | 2 | 1 | 2 | 1 | 2 | 1 | 2 | 1 | 2 | 1 | 2 | 1 | 2 | 1 | 2 |
| 31 | 1 | 2 | 3 | 1 | 2 | 3 | 1 | 2 | 3 | 1 | 2 | 3 | 1 | 2 | 3 | 1 | 2 | 3 | 1 | 2 | 3 | 1 | 2 | 3 | 1 | 2 | 3 | 1 | 2 | 3 |
| 32 | 1 | 2 | 1 | 2 | 1 | 2 | 1 | 2 | 1 | 2 | 1 | 2 | 3 | 1 | 2 | 1 | 2 | 3 | 3 | 3 | 3 | 3 | 3 | 3 | 3 | 3 | 3 | 3 | 3 | 3 |
| 33 | 1 | 2 | 1 | 1 | 2 | 1 | 2 | 1 | 1 | 1 | 2 | 1 | 2 | 1 | 1 | 2 | 1 | 2 | 2 | 1 | 1 | 1 | 1 | 1 | 2 | 3 | 1 | 1 | 2 | 2 |
| 34 | 1 | 1 | 1 | 1 | 2 | 1 | 1 | 1 | 1 | 1 | 1 | 1 | 1 | 2 | 1 | 1 | 1 | 1 | 1 | 1 | 1 | 1 | 1 | 1 | 1 | 2 | 2 | 1 | 1 | 1 |
| 35 | 1 | 1 | 2 | 1 | 2 | 1 | 1 | 1 | 1 | 2 | 1 | 1 | 1 | 2 | 1 | 1 | 2 | 1 | 1 | 3 | 1 | 1 | 1 | 1 | 1 | 1 | 1 | 1 | 1 | 1 |
| 36 | 2 | 1 | 2 | 1 | 1 | 2 | 1 | 1 | 1 | 1 | 1 | 1 | 2 | 1 | 2 | 1 | 2 | 2 | 1 | 1 | 3 | 2 | 1 | 2 | 2 | 2 | 1 | 2 | 1 | 1 |
| 37 | 1 | 1 | 1 | 1 | 1 | 1 | 2 | 1 | 2 | 1 | 2 | 2 | 2 | 1 | 1 | 1 | 2 | 2 | 1 | 2 | 1 | 2 | 1 | 2 | 2 | 1 | 2 | 1 | 3 | 2 |
| 38 | 1 | 2 | 1 | 1 | 2 | 1 | 2 | 2 | 2 | 1 | 2 | 1 | 2 | 2 | 1 | 1 | 1 | 2 | 1 | 1 | 2 | 2 | 2 | 2 | 1 | 1 | 2 | 1 | 3 | 2 |
| 39 | 2 | 2 | 1 | 2 | 1 | 2 | 1 | 2 | 1 | 1 | 1 | 1 | 1 | 2 | 1 | 2 | 2 | 2 | 1 | 2 | 2 | 1 | 1 | 1 | 1 | 1 | 2 | 2 | 1 | 1 |
| 40 | 2 | 2 | 1 | 1 | 1 | 1 | 1 | 1 | 1 | 1 | 2 | 2 | 1 | 1 | 1 | 1 | 2 | 1 | 1 | 1 | 1 | 1 | 1 | 3 | 1 | 1 | 1 | 1 | 1 | 1 |
| 41 | 1 | 1 | 2 | 2 | 2 | 2 | 1 | 1 | 1 | 2 | 1 | 2 | 1 | 1 | 1 | 1 | 2 | 2 | 1 | 2 | 2 | 1 | 1 | 1 | 1 | 2 | 1 | 1 | 1 | 2 |
| 42 | 1 | 1 | 1 | 2 | 1 | 1 | 2 | 2 | 1 | 1 | 2 | 1 | 1 | 2 | 1 | 1 | 1 | 1 | 2 | 1 | 2 | 2 | 1 | 2 | 1 | 1 | 2 | 1 | 2 | 1 |
| 43 | 1 | 2 | 1 | 1 | 1 | 1 | 1 | 2 | 1 | 2 | 1 | 1 | 1 | 1 | 1 | 2 | 1 | 1 | 2 | 1 | 1 | 1 | 1 | 1 | 1 | 2 | 2 | 2 | 2 | 1 |
| 44 | 1 | 2 | 1 | 1 | 1 | 1 | 2 | 1 | 2 | 2 | 1 | 2 | 1 | 1 | 1 | 2 | 1 | 2 | 2 | 1 | 2 | 1 | 1 | 1 | 1 | 1 | 3 | 2 | 1 | 2 |
| 45 | 2 | 2 | 2 | 1 | 1 | 2 | 1 | 2 | 1 | 2 | 1 | 1 | 1 | 1 | 2 | 2 | 1 | 1 | 1 | 2 | 1 | 1 | 1 | 1 | 3 | 2 | 1 | 1 | 1 | 2 |
| 46 | 2 | 2 | 1 | 1 | 2 | 2 | 1 | 1 | 1 | 1 | 1 | 1 | 1 | 1 | 1 | 2 | 1 | 1 | 2 | 2 | 1 | 1 | 1 | 2 | 2 | 2 | 2 | 1 | 1 | 3 |
| 47 | 1 | 2 | 2 | 1 | 2 | 2 | 2 | 2 | 1 | 1 | 1 | 2 | 2 | 1 | 2 | 1 | 1 | 2 | 2 | 1 | 2 | 1 | 1 | 1 | 1 | 1 | 1 | 1 | 2 | 1 |
| 48 | 2 | 1 | 2 | 2 | 2 | 2 | 1 | 2 | 1 | 1 | 2 | 2 | 2 | 1 | 1 | 2 | 1 | 1 | 1 | 1 | 2 | 1 | 2 | 2 | 1 | 2 | 1 | 2 | 1 | 1 |
| 49 | 2 | 1 | 1 | 1 | 1 | 1 | 1 | 1 | 2 | 2 | 1 | 2 | 1 | 2 | 1 | 2 | 1 | 1 | 2 | 1 | 1 | 1 | 1 | 2 | 1 | 1 | 3 | 2 | 2 | 2 |
| 50 | 2 | 1 | 2 | 2 | 1 | 1 | 1 | 1 | 1 | 2 | 1 | 1 | 1 | 1 | 1 | 1 | 1 | 1 | 1 | 2 | 2 | 1 | 2 | 2 | 2 | 2 | 1 | 1 | 1 | 2 |
| 51 | 2 | 2 | 2 | 1 | 1 | 1 | 2 | 2 | 2 | 1 | 2 | 1 | 2 | 2 | 1 | 1 | 1 | 1 | 1 | 1 | 2 | 1 | 2 | 1 | 1 | 1 | 1 | 2 | 1 | 1 |
| 52 | 1 | 1 | 2 | 1 | 2 | 1 | 1 | 1 | 1 | 2 | 1 | 1 | 1 | 2 | 1 | 1 | 2 | 1 | 1 | 2 | 1 | 1 | 2 | 1 | 1 | 3 | 2 | 2 | 2 | 2 |
| 53 | 2 | 2 | 1 | 3 | 2 | 2 | 2 | 2 | 2 | 2 | 2 | 2 | 1 | 1 | 1 | 1 | 1 | 1 | 1 | 1 | 1 | 1 | 1 | 1 | 1 | 1 | 1 | 1 | 1 | 1 |
| 54 | 2 | 1 | 1 | 2 | 2 | 1 | 2 | 1 | 2 | 2 | 2 | 1 | 1 | 1 | 1 | 1 | 2 | 2 | 1 | 1 | 1 | 1 | 1 | 2 | 2 | 2 | 1 | 1 | 1 | 1 |
| 55 | 1 | 1 | 1 | 1 | 1 | 1 | 1 | 1 | 2 | 1 | 1 | 1 | 2 | 1 | 1 | 2 | 1 | 1 | 1 | 2 | 1 | 1 | 2 | 2 | 2 | 1 | 1 | 1 | 2 | 1 |
| 56 | 1 | 2 | 2 | 2 | 2 | 2 | 1 | 1 | 1 | 1 | 2 | 2 | 1 | 1 | 1 | 2 | 1 | 1 | 1 | 1 | 2 | 1 | 1 | 2 | 1 | 1 | 1 | 1 | 3 | 2 |
| 57 | 2 | 1 | 1 | 2 | 1 | 2 | 1 | 1 | 2 | 2 | 1 | 1 | 1 | 2 | 1 | 2 | 1 | 2 | 1 | 1 | 1 | 1 | 2 | 1 | 1 | 1 | 2 | 1 | 3 | 2 |
| 58 | 1 | 1 | 1 | 1 | 1 | 1 | 2 | 1 | 1 | 1 | 2 | 2 | 1 | 1 | 2 | 1 | 1 | 2 | 1 | 2 | 2 | 1 | 2 | 1 | 1 | 2 | 1 | 1 | 1 | 2 |
| 69 | 1 | 2 | 1 | 2 | 1 | 2 | 2 | 2 | 1 | 2 | 1 | 2 | 1 | 2 | 2 | 1 | 1 | 1 | 1 | 1 | 2 | 2 | 2 | 1 | 2 | 1 | 2 | 1 | 1 | 1 |
| 60 | 1 | 1 | 2 | 1 | 2 | 2 | 1 | 2 | 1 | 1 | 1 | 1 | 1 | 1 | 1 | 1 | 1 | 1 | 1 | 1 | 1 | 1 | 1 | 1 | 2 | 1 | 2 | 2 | 1 | 2 |
| 61 | 1 | 1 | 2 | 2 | 2 | 2 | 1 | 1 | 1 | 2 | 2 | 2 | 2 | 1 | 1 | 1 | 2 | 1 | 1 | 1 | 1 | 2 | 1 | 2 | 1 | 1 | 2 | 1 | 2 | 1 |
| 62 | 1 | 1 | 1 | 1 | 1 | 1 | 1 | 1 | 1 | 1 | 1 | 1 | 1 | 1 | 1 | 1 | 1 | 1 | 1 | 1 | 1 | 1 | 1 | 1 | 1 | 1 | 1 | 1 | 1 | 1 |
| 63 | 1 | 1 | 1 | 1 | 1 | 1 | 1 | 1 | 1 | 1 | 1 | 1 | 1 | 1 | 1 | 1 | 1 | 1 | 1 | 1 | 1 | 1 | 3 | 2 | 2 | 2 | 2 | 1 | 1 | 1 |
| 64 | 1 | 1 | 1 | 1 | 1 | 1 | 1 | 1 | 1 | 1 | 1 | 1 | 1 | 1 | 1 | 1 | 1 | 1 | 2 | 1 | 1 | 1 | 1 | 1 | 1 | 1 | 1 | 1 | 1 | 1 |
| 65 | 3 | 3 | 2 | 3 | 1 | 2 | 2 | 2 | 3 | 2 | 3 | 1 | 2 | 3 | 1 | 3 | 3 | 2 | 3 | 3 | 1 | 1 | 2 | 3 | 3 | 2 | 3 | 3 | 2 | 1 |
| 66 | 1 | 2 | 1 | 2 | 2 | 2 | 1 | 2 | 2 | 3 | 3 | 2 | 3 | 3 | 2 | 1 | 3 | 2 | 2 | 1 | 2 | 3 | 1 | 2 | 1 | 3 | 1 | 1 | 2 | 1 |
| 67 | 2 | 2 | 2 | 1 | 3 | 2 | 3 | 3 | 1 | 1 | 1 | 2 | 2 | 2 | 3 | 1 | 1 | 3 | 2 | 2 | 2 | 2 | 2 | 3 | 1 | 2 | 2 | 1 | 2 | 3 |
| 68 | 3 | 2 | 2 | 1 | 2 | 3 | 1 | 1 | 1 | 3 | 3 | 1 | 1 | 3 | 2 | 1 | 1 | 1 | 2 | 1 | 3 | 3 | 3 | 3 | 2 | 1 | 3 | 2 | 1 | 1 |
| 69 | 1 | 3 | 2 | 3 | 1 | 2 | 3 | 2 | 1 | 3 | 2 | 1 | 3 | 3 | 1 | 2 | 1 | 1 | 1 | 3 | 1 | 2 | 2 | 1 | 1 | 1 | 2 | 1 | 3 | 2 |
| 70 | 2 | 3 | 3 | 2 | 1 | 1 | 1 | 3 | 3 | 2 | 1 | 1 | 1 | 1 | 1 | 2 | 2 | 1 | 1 | 2 | 2 | 1 | 2 | 1 | 3 | 3 | 1 | 2 | 1 | 1 |
| 71 | 3 | 1 | 3 | 2 | 2 | 2 | 3 | 3 | 1 | 1 | 1 | 1 | 2 | 1 | 2 | 1 | 1 | 2 | 3 | 3 | 3 | 2 | 1 | 3 | 2 | 2 | 3 | 1 | 2 | 1 |
| 72 | 3 | 2 | 3 | 3 | 2 | 2 | 1 | 2 | 2 | 2 | 2 | 3 | 1 | 2 | 1 | 3 | 1 | 2 | 1 | 2 | 1 | 1 | 3 | 3 | 3 | 2 | 2 | 2 | 3 | 1 |
| 73 | 3 | 1 | 3 | 1 | 3 | 3 | 1 | 2 | 3 | 2 | 3 | 2 | 2 | 2 | 3 | 1 | 2 | 1 | 2 | 3 | 1 | 1 | 2 | 2 | 2 | 2 | 2 | 1 | 1 | 1 |
| 74 | 1 | 3 | 3 | 1 | 3 | 3 | 3 | 1 | 3 | 3 | 1 | 3 | 3 | 1 | 2 | 3 | 3 | 1 | 2 | 2 | 1 | 1 | 1 | 1 | 2 | 1 | 2 | 3 | 1 | 2 |
| 75 | 3 | 3 | 2 | 2 | 2 | 2 | 3 | 1 | 2 | 3 | 2 | 2 | 2 | 2 | 1 | 1 | 1 | 2 | 3 | 3 | 3 | 1 | 2 | 2 | 1 | 3 | 2 | 3 | 1 | 3 |
| 76 | 2 | 1 | 1 | 2 | 1 | 3 | 3 | 1 | 1 | 3 | 1 | 1 | 2 | 2 | 2 | 3 | 3 | 1 | 1 | 2 | 1 | 2 | 1 | 3 | 2 | 2 | 2 | 2 | 3 | 2 |
| 77 | 3 | 3 | 3 | 1 | 3 | 2 | 1 | 2 | 2 | 3 | 1 | 2 | 2 | 1 | 2 | 3 | 1 | 1 | 2 | 2 | 2 | 2 | 3 | 2 | 2 | 1 | 3 | 1 | 3 | 3 |
| 78 | 2 | 1 | 2 | 2 | 3 | 1 | 3 | 2 | 3 | 2 | 2 | 3 | 2 | 1 | 3 | 3 | 2 | 1 | 2 | 2 | 1 | 3 | 3 | 3 | 2 | 3 | 1 | 2 | 1 | 2 |
| 79 | 2 | 3 | 1 | 1 | 3 | 2 | 2 | 2 | 1 | 3 | 1 | 2 | 2 | 2 | 2 | 1 | 1 | 3 | 3 | 3 | 2 | 1 | 2 | 3 | 1 | 2 | 2 | 2 | 1 | 2 |
| 80 | 1 | 1 | 1 | 2 | 3 | 2 | 2 | 3 | 3 | 2 | 1 | 3 | 2 | 3 | 3 | 2 | 3 | 2 | 2 | 2 | 3 | 3 | 2 | 1 | 2 | 2 | 1 | 3 | 2 | 3 |

Sequence Position (positions 31–60)

| Sequence ID | 31 | 32 | 33 | 34 | 35 | 36 | 37 | 38 | 39 | 40 | 41 | 42 | 43 | 44 | 45 | 46 | 47 | 48 | 49 | 50 | 51 | 52 | 53 | 54 | 55 | 56 | 57 | 58 | 59 | 60 |
|---|---|---|---|---|---|---|---|---|---|---|---|---|---|---|---|---|---|---|---|---|---|---|---|---|---|---|---|---|---|---|
| 33 | 2 | 3 | 1 | 2 | 1 | 1 | 2 | 1 | 2 | 2 | 1 | 1 | 1 | 2 | 1 | 1 | 2 | 2 | 2 | 2 | 2 | 2 | 2 | 1 | 1 | 1 | 2 | 2 | 1 | 3 |
| 34 | 1 | 3 | 1 | 3 | 2 | 1 | 1 | 1 | 1 | 2 | 2 | 2 | 2 | 2 | 1 | 1 | 1 | 1 | 1 | 1 | 1 | 1 | 1 | 1 | 2 | 1 | 1 | 2 | 3 | 1 |
| 35 | 1 | 1 | 1 | 1 | 2 | 1 | 1 | 1 | 1 | 1 | 2 | 1 | 2 | 3 | 2 | 2 | 2 | 1 | 1 | 1 | 2 | 1 | 1 | 1 | 1 | 1 | 1 | 2 | 1 | 3 |
| 36 | 1 | 1 | 2 | 1 | 1 | 2 | 2 | 1 | 2 | 2 | 1 | 2 | 2 | 1 | 2 | 1 | 2 | 2 | 1 | 1 | 1 | 2 | 3 | 2 | 1 | 2 | 1 | 2 | 3 | 2 |
| 37 | 1 | 1 | 2 | 1 | 1 | 1 | 2 | 1 | 2 | 2 | 1 | 1 | 1 | 2 | 3 | 2 | 1 | 2 | 2 | 1 | 1 | 1 | 2 | 2 | 2 | 2 | 1 | 2 | 3 | 1 |
| 38 | 1 | 1 | 1 | 2 | 1 | 1 | 1 | 1 | 1 | 2 | 2 | 2 | 1 | 2 | 1 | 1 | 1 | 1 | 1 | 1 | 1 | 1 | 1 | 1 | 2 | 2 | 2 | 3 | 3 | 2 |
| 39 | 3 | 1 | 1 | 2 | 1 | 1 | 2 | 1 | 2 | 1 | 1 | 2 | 1 | 2 | 3 | 1 | 1 | 2 | 1 | 1 | 2 | 1 | 1 | 1 | 2 | 1 | 1 | 2 | 3 | 2 |
| 40 | 1 | 1 | 2 | 3 | 1 | 1 | 1 | 1 | 1 | 1 | 1 | 1 | 1 | 1 | 1 | 1 | 1 | 1 | 1 | 1 | 1 | 1 | 1 | 1 | 1 | 1 | 1 | 1 | 3 | 1 |
| 41 | 2 | 1 | 1 | 1 | 2 | 3 | 1 | 2 | 2 | 1 | 3 | 2 | 1 | 2 | 1 | 1 | 2 | 1 | 1 | 2 | 2 | 2 | 1 | 2 | 2 | 2 | 2 | 1 | 3 | 2 |
| 42 | 1 | 1 | 1 | 1 | 1 | 2 | 2 | 1 | 3 | 3 | 1 | 1 | 2 | 1 | 2 | 1 | 2 | 1 | 1 | 2 | 1 | 3 | 1 | 1 | 1 | 3 | 1 | 1 | 2 | 3 |
| 43 | 2 | 1 | 2 | 1 | 1 | 1 | 1 | 1 | 3 | 3 | 2 | 2 | 1 | 3 | 2 | 2 | 2 | 2 | 1 | 1 | 2 | 3 | 1 | 1 | 1 | 1 | 1 | 1 | 1 | 2 |
| 44 | 2 | 1 | 2 | 2 | 1 | 1 | 2 | 1 | 1 | 1 | 1 | 1 | 1 | 1 | 1 | 1 | 2 | 1 | 1 | 1 | 2 | 2 | 3 | 1 | 1 | 1 | 1 | 2 | 1 | 3 |
| 45 | 1 | 1 | 1 | 1 | 1 | 1 | 1 | 1 | 1 | 1 | 2 | 2 | 1 | 1 | 2 | 1 | 1 | 2 | 2 | 2 | 1 | 1 | 2 | 1 | 1 | 1 | 2 | 3 | 1 | 3 |
| 46 | 2 | 2 | 2 | 2 | 2 | 2 | 2 | 2 | 3 | 1 | 2 | 1 | 1 | 1 | 1 | 1 | 2 | 1 | 1 | 2 | 1 | 1 | 1 | 1 | 2 | 1 | 1 | 1 | 3 | 2 |
| 47 | 1 | 3 | 2 | 2 | 2 | 2 | 2 | 2 | 3 | 1 | 1 | 1 | 1 | 2 | 1 | 1 | 2 | 2 | 1 | 1 | 1 | 2 | 2 | 2 | 2 | 1 | 2 | 1 | 3 | 2 |
| 48 | 1 | 1 | 1 | 3 | 2 | 2 | 2 | 2 | 2 | 3 | 1 | 1 | 2 | 2 | 1 | 1 | 1 | 1 | 2 | 1 | 2 | 1 | 2 | 2 | 2 | 2 | 1 | 1 | 3 | 1 |
| 49 | 2 | 2 | 2 | 2 | 2 | 2 | 2 | 2 | 2 | 2 | 2 | 2 | 2 | 2 | 2 | 2 | 3 | 1 | 1 | 1 | 2 | 1 | 1 | 1 | 1 | 1 | 1 | 1 | 3 | 2 |
| 50 | 2 | 2 | 1 | 1 | 1 | 3 | 2 | 2 | 2 | 1 | 1 | 1 | 2 | 1 | 1 | 1 | 1 | 2 | 3 | 1 | 2 | 2 | 1 | 1 | 1 | 2 | 1 | 2 | 3 | 2 |
| 51 | 3 | 2 | 2 | 2 | 2 | 2 | 2 | 2 | 2 | 2 | 2 | 2 | 2 | 2 | 2 | 3 | 1 | 2 | 1 | 1 | 2 | 1 | 1 | 2 | 1 | 1 | 1 | 1 | 3 | 1 |
| 52 | 2 | 2 | 2 | 2 | 2 | 2 | 2 | 2 | 2 | 2 | 2 | 2 | 2 | 2 | 2 | 2 | 2 | 2 | 2 | 2 | 2 | 2 | 2 | 2 | 2 | 3 | 1 | 1 | 3 | 1 |
| 53 | 1 | 1 | 1 | 1 | 1 | 1 | 1 | 1 | 1 | 1 | 1 | 1 | 1 | 1 | 1 | 1 | 1 | 1 | 1 | 3 | 1 | 1 | 1 | 1 | 1 | 2 | 1 | 2 | 3 | 1 |
| 54 | 3 | 2 | 2 | 2 | 2 | 2 | 2 | 2 | 2 | 2 | 2 | 2 | 2 | 2 | 2 | 2 | 2 | 2 | 2 | 2 | 2 | 2 | 2 | 2 | 3 | 1 | 2 | 1 | 3 | 2 |
| 55 | 1 | 2 | 2 | 2 | 2 | 2 | 1 | 1 | 1 | 1 | 1 | 2 | 2 | 1 | 2 | 1 | 1 | 2 | 2 | 2 | 1 | 1 | 1 | 2 | 3 | 1 | 1 | 1 | 1 | 3 |
| 56 | 1 | 2 | 1 | 1 | 1 | 2 | 2 | 2 | 2 | 1 | 1 | 1 | 1 | 1 | 1 | 1 | 1 | 2 | 1 | 2 | 3 | 1 | 1 | 1 | 1 | 1 | 1 | 1 | 3 | 1 |
| 57 | 1 | 1 | 1 | 2 | 1 | 2 | 1 | 1 | 1 | 1 | 1 | 2 | 1 | 1 | 1 | 1 | 1 | 1 | 1 | 1 | 1 | 1 | 1 | 3 | 2 | 2 | 2 | 2 | 3 | 2 |
| 58 | 1 | 3 | 2 | 1 | 2 | 1 | 1 | 1 | 2 | 1 | 1 | 1 | 2 | 1 | 2 | 2 | 1 | 1 | 2 | 1 | 1 | 1 | 1 | 2 | 3 | 1 | 1 | 2 | 1 | 3 |
| 69 | 1 | 2 | 1 | 1 | 1 | 1 | 1 | 1 | 1 | 1 | 1 | 1 | 1 | 1 | 1 | 1 | 1 | 2 | 2 | 1 | 1 | 2 | 3 | 1 | 1 | 1 | 3 | 1 | 3 | 1 |
| 60 | 1 | 2 | 1 | 1 | 2 | 1 | 1 | 1 | 2 | 2 | 1 | 1 | 1 | 1 | 1 | 2 | 1 | 1 | 1 | 2 | 2 | 2 | 2 | 1 | 3 | 1 | 3 | 1 | 2 | 3 |
| 61 | 2 | 2 | 1 | 1 | 2 | 2 | 2 | 2 | 2 | 2 | 1 | 2 | 2 | 2 | 2 | 2 | 2 | 2 | 2 | 2 | 2 | 2 | 2 | 2 | 2 | 2 | 2 | 2 | 3 | 2 |
| 62 | 1 | 1 | 1 | 1 | 1 | 1 | 1 | 2 | 2 | 2 | 1 | 2 | 1 | 1 | 1 | 1 | 3 | 2 | 2 | 2 | 2 | 2 | 2 | 1 | 3 | 1 | 1 | 2 | 3 | 2 |
| 63 | 2 | 2 | 1 | 2 | 1 | 2 | 2 | 2 | 1 | 1 | 2 | 2 | 1 | 2 | 1 | 1 | 1 | 1 | 1 | 1 | 1 | 1 | 2 | 2 | 2 | 1 | 3 | 2 | 2 | 3 |
| 64 | 1 | 1 | 1 | 1 | 1 | 1 | 1 | 1 | 1 | 1 | 2 | 1 | 1 | 1 | 1 | 1 | 1 | 1 | 1 | 1 | 3 | 1 | 3 | 1 | 1 | 1 | 1 | 1 | 3 | 2 |
| 65 | 2 | 3 | 2 | 2 | 1 | 2 | 1 | 1 | 2 | 2 | 2 | 1 | 2 | 1 | 1 | 2 | 2 | 1 | 2 | 1 | 2 | 1 | 1 | 2 | 1 | 1 | 1 | 1 | 3 | 1 |
| 66 | 2 | 1 | 3 | 3 | 2 | 3 | 3 | 2 | 1 | 1 | 3 | 1 | 2 | 1 | 2 | 3 | 2 | 3 | 3 | 3 | 1 | 3 | 3 | 3 | 2 | 2 | 2 | 3 | 3 | 1 |
| 67 | 1 | 1 | 1 | 2 | 1 | 3 | 2 | 1 | 1 | 3 | 2 | 2 | 2 | 3 | 1 | 2 | 1 | 3 | 1 | 1 | 1 | 1 | 3 | 1 | 3 | 3 | 1 | 1 | 2 | 1 |
| 68 | 3 | 1 | 1 | 2 | 1 | 2 | 2 | 2 | 1 | 2 | 3 | 1 | 2 | 2 | 2 | 1 | 3 | 3 | 2 | 3 | 1 | 2 | 1 | 3 | 2 | 1 | 1 | 3 | 1 | 3 |
| 69 | 3 | 2 | 2 | 1 | 1 | 1 | 1 | 2 | 3 | 3 | 2 | 1 | 2 | 2 | 3 | 1 | 1 | 3 | 3 | 2 | 1 | 2 | 3 | 1 | 3 | 3 | 1 | 1 | 2 | 2 |
| 70 | 1 | 3 | 2 | 3 | 2 | 2 | 3 | 1 | 2 | 2 | 1 | 2 | 3 | 3 | 3 | 3 | 1 | 1 | 3 | 3 | 2 | 3 | 1 | 3 | 1 | 3 | 3 | 2 | 3 | 1 |
| 71 | 2 | 1 | 1 | 2 | 1 | 3 | 2 | 3 | 3 | 2 | 1 | 3 | 1 | 3 | 2 | 3 | 2 | 3 | 1 | 3 | 1 | 1 | 2 | 2 | 2 | 1 | 2 | 1 | 3 | 2 |
| 72 | 1 | 3 | 2 | 2 | 1 | 1 | 3 | 3 | 1 | 3 | 2 | 2 | 2 | 2 | 1 | 3 | 3 | 3 | 2 | 1 | 2 | 1 | 1 | 3 | 1 | 3 | 3 | 2 | 1 | 3 |
| 73 | 1 | 2 | 3 | 3 | 2 | 2 | 1 | 1 | 1 | 2 | 3 | 1 | 2 | 1 | 2 | 1 | 1 | 2 | 3 | 1 | 1 | 3 | 1 | 3 | 3 | 3 | 1 | 2 | 2 | 1 |
| 74 | 1 | 3 | 2 | 3 | 1 | 3 | 2 | 2 | 3 | 3 | 3 | 3 | 2 | 2 | 2 | 1 | 2 | 3 | 1 | 1 | 1 | 3 | 2 | 3 | 2 | 1 | 3 | 2 | 3 | 3 |
| 75 | 3 | 2 | 1 | 3 | 3 | 2 | 1 | 1 | 3 | 2 | 1 | 2 | 1 | 3 | 3 | 1 | 2 | 3 | 1 | 2 | 1 | 3 | 3 | 3 | 3 | 2 | 3 | 2 | 3 | 3 |
| 76 | 1 | 3 | 2 | 2 | 3 | 1 | 2 | 2 | 2 | 1 | 1 | 1 | 2 | 2 | 2 | 1 | 3 | 1 | 2 | 2 | 2 | 1 | 2 | 2 | 3 | 1 | 3 | 2 | 3 | 3 |
| 77 | 3 | 1 | 1 | 1 | 3 | 3 | 3 | 2 | 2 | 1 | 1 | 3 | 2 | 3 | 1 | 3 | 2 | 2 | 3 | 2 | 1 | 2 | 1 | 1 | 2 | 2 | 3 | 2 | 3 | 2 |
| 78 | 3 | 2 | 1 | 1 | 1 | 3 | 2 | 1 | 3 | 3 | 3 | 1 | 1 | 3 | 3 | 2 | 2 | 1 | 1 | 2 | 1 | 2 | 1 | 2 | 2 | 2 | 2 | 3 | 2 | 3 |
| 79 | 1 | 2 | 1 | 2 | 2 | 3 | 2 | 1 | 2 | 2 | 1 | 3 | 3 | 2 | 2 | 2 | 3 | 2 | 1 | 1 | 3 | 3 | 3 | 3 | 2 | 1 | 3 | 1 | 3 | 2 |
| 80 | 2 | 3 | 1 | 1 | 1 | 1 | 2 | 2 | 3 | 2 | 1 | 1 | 2 | 1 | 1 | 3 | 2 | 2 | 1 | 3 | 2 | 1 | 1 | 2 | 3 | 1 | 2 | 2 | 1 | 2 |

#### Appendix 2: Unigram Predictive-Looking on Object’s First Appearance GAM Visualization

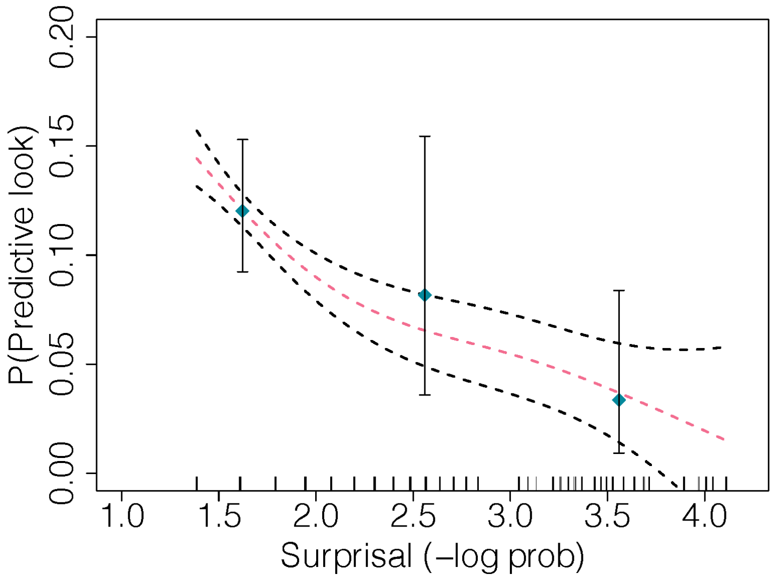


Figure 3: Probability of Predictive-Look on Object’s First Appearance as a Function of Unigram Surprisal. Subjects’ probability of predictively looking to the currently active object (y-axis) on their first appearance as a function of surprisal (x-axis) as measured by the unigram model; the smooth curve shows the fit of a generalized additive model with 95% confidence interval.

### Appendix 3: Transitional Look-Away GAM Visualization

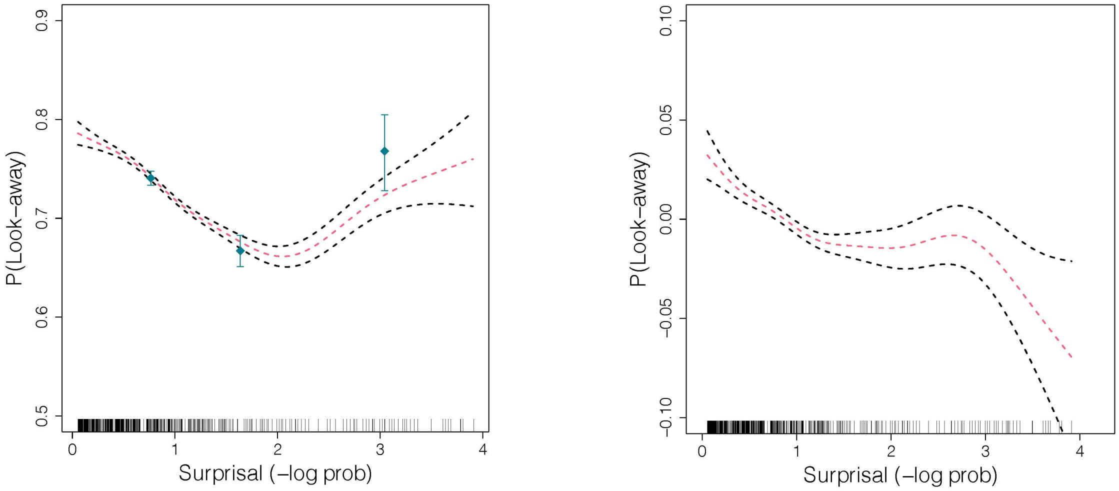


Fig. 4. Look-Away Probability as a Function of Bigram Surprisal. (a) Subjects' probability of looking away (y-axis) as a function of surprisal (x-axis) as measured by the transitional  model. The points and error bars show the raw probability of looking away; the smooth curve shows the fit of a generalized additive model with standard errors. Vertical tick marks show values of surprisal attained in the experiment. (b) The relationship between look-away probability (y–axis) and bigram surprisal (x-axis), while controlling for all covariate factors.

### Appendix 4: Transitional Reaction Time GAM Visualization

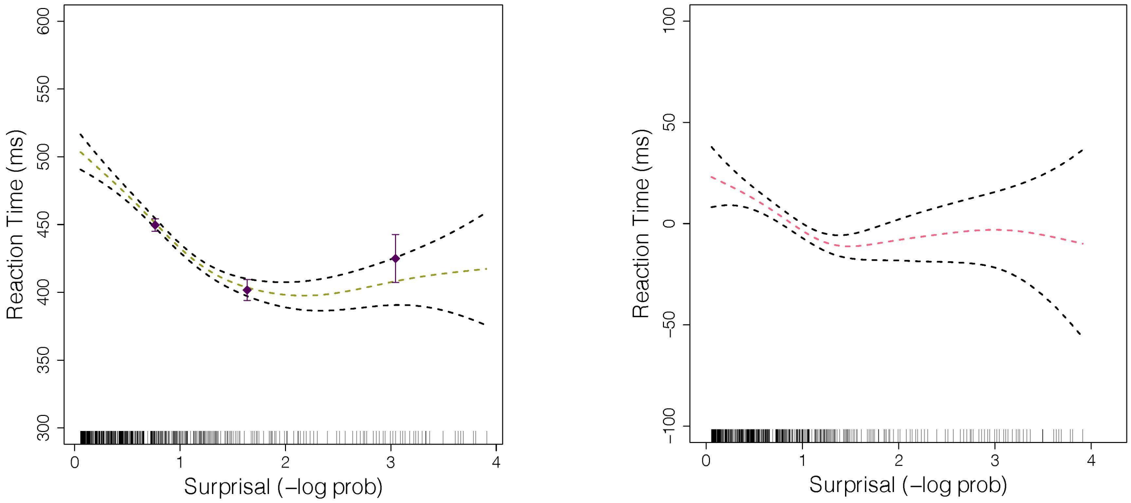


Fig. 5. Reaction Time (ms) as a Function of Bigram Surprisal. (a) Subjects' reaction time (latency) to fixate the active object (*y*-axis) as a function of surprisal (*x*-axis) as measured by the transitional model; the smooth curve shows the fit of a generalized additive model with standard errors. *(b)* Reaction Time (*y*–axis) and bigram surprisal (*x*-axis), while controlling for all factors.

#### Appendix 5: Transitional Predictive-Looking on Object's First Appearance GAM Visualization

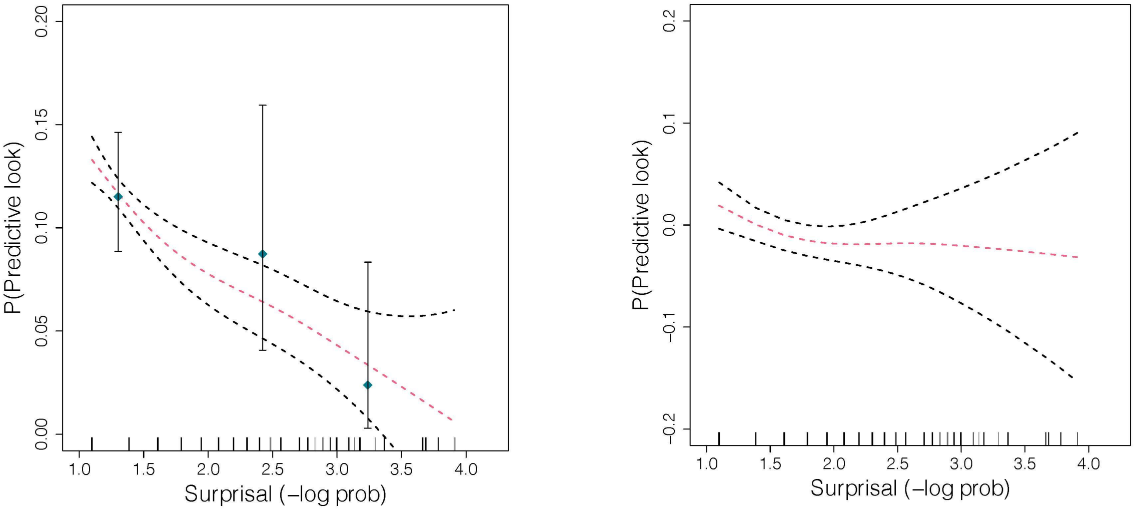


Fig. 6. Probability of Predictive-Look on Object's First Appearance as a Function of Bigram Surprisal. (a) Subjects' probability of predictively looking to the currently active object (y-axis) on their first appearance as a function of surprisal (x-axis) as measured by the transitional model; the smooth curve shows the fit of a generalized additive model with standard errors.(b) Subjects' probability of predictively looking to the currently active object (y-axis) on their first appearance as a function of bigram surprisal (x-axis), while controlling for all factors.

#### Appendix 6: Markov Dirichlet-Multinomial Model (Ideal Learner Model)

Intuitively, learners observe how many times each event occurs in the world, and then use these event counts to infer an underlying probability model of their observations. In the experiment, there are three possible events corresponding to which of three objects appears from behind its box.

An observer who sees only a single event happen would not likely infer that the single observed event is the only one possible (i.e., has probability of 1); Instead, observers likely bring expectations to this learning task. In the MDM model used here, this prior expectation is parameterized by a single free parameter, $\alpha$, which controls the strength of the learner's prior belief that the distribution of events is uniform. As $\alpha$ gets large, the model has strong prior beliefs that the distribution of events in the world is uniform; as $\alpha$ approaches zero, the model believes more strongly that the true distribution closely resembles that of the empirically observed event counts. In modeling, we chose a value of $\alpha = 1$, corresponding to a uniform prior expectation about the distribution of events. However, the qualitative results–in particular, the U-shaped relationship between surprisal and look-away probability–do not depend strongly on the choice of $\alpha$.

Formally, suppose there are $N$events, $x_1, x_2, \ldots, x_N$and the $i$th event has been observed $c_i$ times. We are interested in estimating (or scoring) a multinomial distribution parameterized by $\theta = \left(\theta_1, \theta_2, \ldots, \theta_N\right)$where $\theta_i$ is the true (unobserved) probability of event $x_i$. Under a Dirichlet-Multinomial model,

(1)
$$P\left(\theta \mid c_1, \ldots c_N, \alpha\right) = \frac{1}{B} \prod_{i=1}^{N} \theta_i^{\alpha + c_i - 1}$$

where $B$ is a normalizing constant that depends on the $c_i$ and $\alpha$. That is, after observing each event type occur some number of times, the infant may form a representation, $\theta$, of their guess at the true distribution of events. Every distribution can be scored according to Equation 1, allowing one to compute how strongly a learner should believe that any particular $\theta$ is the correct one. We predict that macaques' reaction times to the pop-up objects will depend upon the surprisal of that current event, which is determined by both the previously observed events and the identity of the current event. We predict that events of either very low surprisal (highly predictable) or very high surprisal (highly unexpected) will be more likely to trigger a shorter latency than events with moderate surprisal.

When the $i$th event occurs, the main variable of interest here is its negative log probability according to the model. We compute this by integrating over the above posterior distribution on $\theta$. This corresponds to a measure of the information conveyed by observing event $i$ according to an ideal Bayesian learner who had seen all previous events. We predicted that macaques would be faster to look at events that contained either too little or too much information, giving a U-shaped (quadratic) relationship between this negative log probability measure and the actual observed look-away probability.

#### Appendix 7: Tables of Regression Results

**Surprisal Term Coefficients for Reaction-Time Regression**

| | | Linear | | Quadratic | | GAM Trend |
|---|---|---|---|---|---|---|
| | | Coef. | P-value | Coef. | P-value | |
| Unigram | RAW | -64.68 | *** | 14.37 | ** | U |
| | CONTROLLED | -26.19 | * | 6.00 | * | U |
| Transitional | RAW | -34.00 | * | 7.63 | ns | Shallow U |
| | CONTROLLED | -10.81 | ns | 3.05 | ns | Flat |
| Significance codes: ‘ *** ’ < 0.001 ‘ ** ’ < 0.01 ‘ * ’ < 0.05 ‘· ’ < 0.1 ‘ ns’ < 1 | | | | | | |

**Surprisal Term Coefficients for Look-Away Regression**

| | | Linear | | Quadratic | | GAM Trend |
|---|---|---|---|---|---|---|
| | | Coef. | P-value | Coef. | P-value | |
| Unigram | RAW | -0.447 | *** | 0.108 | *** | U |
| | CONTROLLED | -0.156 | · | 0.025 | ns | U |
| Transitional | RAW | -0.214 | *** | 0.055 | ** | Shallow U |
| | CONTROLLED | -0.065 | ns | -0.002 | ns | Flat |
| Significance codes: ‘ *** ’ < 0.001 ‘ ** ’ < 0.01 ‘ * ’ < 0.05 ‘· ’ < 0.1 ‘ ns’ < 1 | | | | | | |

**Surprisal Term Coefficients for Predictive-Look with First Appearance Regression**

| | | Linear | | Quadratic | | GAM Trend |
|---|---|---|---|---|---|---|
| | | Coef. | P-value | Coef. | P-value | |
| Unigram | RAW | -0.77 | ns | 0.07 | ns | Decreasing |
| | CONTROLLED | -2.72 | ** | 1.55 | * | Decreasing |
| Transitional | RAW | -0.47 | ns | 0.01 | ns | Decreasing |
| | CONTROLLED | -0.36 | ns | 0.09 | ns | Flat |
| Significance codes: ‘ *** ’ < 0.001 ‘ ** ’ < 0.01 ‘ * ’ < 0.05 ‘· ’ < 0.1 ‘ ns’ < 1 | | | | | | |
